# Extended logistic growth model for heterogeneous populations

**DOI:** 10.1101/231100

**Authors:** Wang Jin, Scott W McCue, Matthew J Simpson

## Abstract

Cell proliferation is the most important cellular-level mechanism responsible for regulating cell population dynamics in living tissues. Modern experimental procedures show that the proliferation rates of individual cells can vary significantly within the same cell line. However, in the mathematical biology literature, cell proliferation is typically modelled using a classical logistic equation which neglects variations in the proliferation rate. In this work, we consider a discrete mathematical model of cell migration and cell proliferation, modulated by volume exclusion (crowding) effects, with variable rates of proliferation across the total population. We refer to this variability as *heterogeneity.* Constructing the continuum limit of the discrete model leads to a generalisation of the classical logistic growth model. Comparing numerical solutions of the model to averaged data from discrete simulations shows that the new model captures the key features of the discrete process. Applying the extended logistic model to simulate a proliferation assay using rates from recent experimental literature shows that neglecting the role of heterogeneity can, at times, lead to misleading results.

## 1 Introduction

Cell proliferation is essential for regulating the dynamics of cell populations, and plays a vital role in collective cell spreading, cancer progression and tissue regeneration (Eladdadi and Isaacson, 2008; Evan and Vousden, 2001; Haridas et al., 2017; Pavlath et al., 1998). While it is clear that cells from different cell lines proliferate at different rates (Hayflick, 1965), recent experimental methods indicate that heterogeneity in cell proliferation arises even within the same cell line (Bajar et al., 2016; Guan et al., 2014; Sakaue-Sawano et al., 2008).

Many different types of experiments are used to quantify cell proliferation (An et al., 2001; Azzarone and Macieira-Coelho, 1982; Haass et al., 2014; Hayflick, 1965; Jin et al., 2017; Kaneoka et al., 1983; Willaime et al., 2013). The complexity of these experiments varies from simple *in vitro* proliferation assays in which the net expansion of a population of cells is observed and measured, such as the experiment shown in Figure 1, to more sophisticated experiments that use fluorescent cell cycle indicators to measure the duration of different phases of the cell cycle for individual cells (Haass et al., 2014; Sakaue-Sawano et al., 2008; Vittadello et al., 2018). A standard measure of cell proliferation is the doubling time, which is a measure of the duration of time required for a population of cells, at low density, to double (Hayflick, 1965; Jin et al., 2016). The doubling time quantifies cell proliferation from the perspective of the entire population, and any kind of variability amongst individual cells in the population is neglected. Modern experimental approaches, such as individual-level fluorescent cell cycle indicators and micro collagen gel arrays, allow us to quantify variations in the cell cycle of individual cells (Guan et al., 2014; Haass et al., 2014). This individual-level data shows that proliferation rates of individual cells can vary significantly within the same cell line (Guan et al., 2014; Haass et al., 2014). Mathematical models are often used to mimic cell biology experiments, and to quantify rates of cell proliferation (Cai et al., 2007; Jin et al., 2017; Nardini et al., 2016). One approach is to apply an individual-level, agent-based model (Frascoli et al, 2014). In this kind of model, agents represent individual cells, and these agents migrate and proliferate according to certain rules thought to be relevant to the application of interest (Treloar et al., 2014). Although agent-based models offer the capability to investigate individual-level details, most of these models adopt a conventional assumption that the rate of proliferation of individual cells in the population is taken to be a constant. This assumption, however, may not be applicable to real situations where the proliferation rate of individual cells in the population varies significantly.

**Fig.1.**
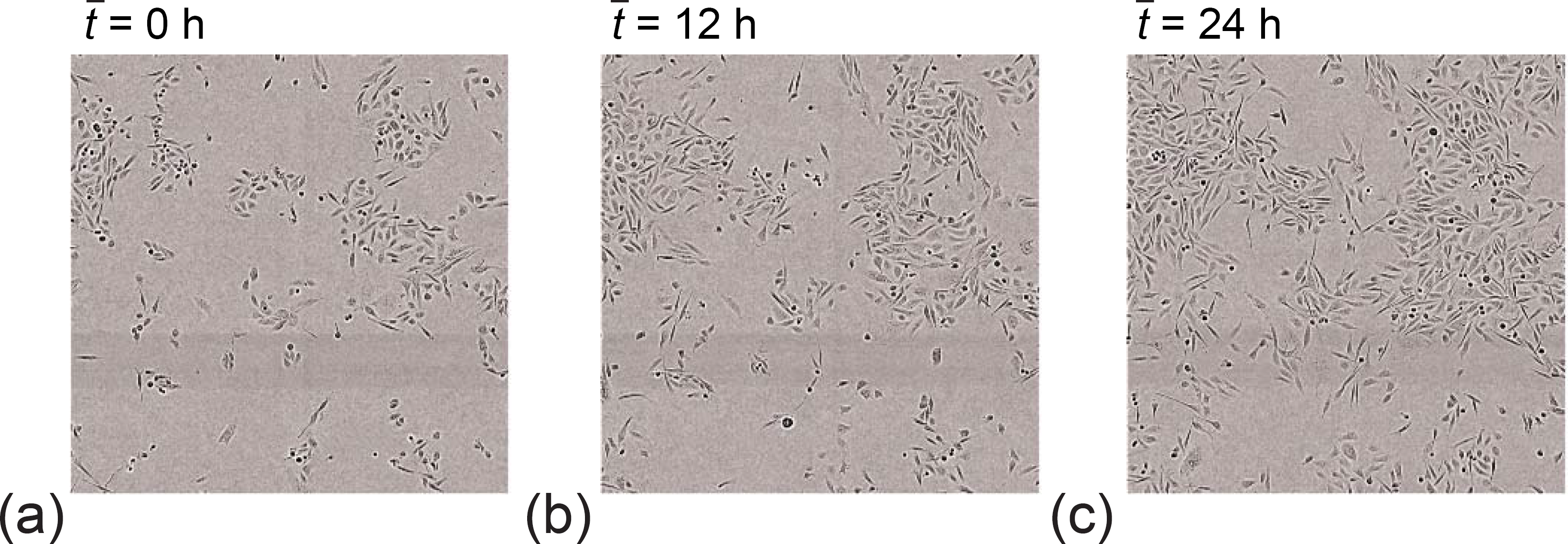
*In vitro* cell proliferation assay. Population of PC-3 prostate cancer cells in a square field of view, of side length 1440 μm. Images correspond to (*a*) 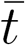 = 0 h, (*b*) 12 h, and (*c*) 24 h (Browning et al., 2018). Reproduced from Browning et al. (2018) with permission.

The most commonly-used continuum model of cell proliferation is the classical logistic growth model (Cai et al., 2007; Jin et al., 2016; Maini et al., 2004; Sen-gers et al., 2007; Sheardown and Cheng, 1996; Sherratt and Murray 1990; Vo et al., 2015; Warne et al. 2017). Although the classical logistic growth model is widely used to estimate the growth rate for populations of cells, there is an increasing awareness in the mathematical biology literature that cell populations do not always grow logistically (e.g. Gerlee, 2013; Powell et al., 2017; Sarapata and de Pillis 2014; Sewalt et al. 2016; West et al. 2001; Neufeld et al. 2017), and generalisations of the logistic growth model have been proposed (Jolicoeur and Pontier, 1989; Tsoularis and Wallace, 2002). Other types of models, where population growth is explicitly coupled to external factors, such as light availability (Pozzobon and Perré, 2018) and interactions with other populations (García-Algarra et al., 2014), have also been developed for specific biological applications. However, a limitation in each of these modelling frameworks is that the cell proliferation rate is treated as a constant, which amounts to neglecting heterogeneity.

In this work we consider a discrete modelling framework in which we deliberately introduce heterogeneity in the rates of migration and proliferation. The continuum limit description of the discrete model leads to a complicated system of reaction-diffusion equations which simplifies to a generalisation of the classical logistic growth model when we apply the model to situations where there are, on average, no spatial variations in the agent density, such as the experimental image in Figure 1. We apply the extended logistic model to simulate a proliferation assay using a distribution of heterogeneous proliferation rates that are estimated from the cell biology literature.

We show that neglecting the role of heterogeneity can produce misleading results when population dynamics are interpreted in a standard way by simply calibrating the solution of the classical logistic growth model to match the data. Comparing the results from the logistic growth model and the extended model illustrates that the logistic growth model does not perform well in some cases. As we show, in these cases where the standard approach fails to capture the growth dynamics of heterogeneous populations, the new extended model performs very well. Unlike the classic logistic growth model, the extended model does not have an exact solution. With this in mind, we provide analytical insight into the role of heterogeneity by constructing approximate perturbation solutions in the limit of small variation in the proliferation rate.

Throughout this study we use a combination of dimensional and dimensionless parameters. Mathematical models, both discrete and continuum, are first presented using dimensional variables and dimensional parameters. All dimensional quantities are indicated with an overbar. Later, when we apply the mathematical models to experimental data, and when we present some analysis of the mathematical models, we always work with dimensionless variables and parameters for which the overbar notation is dropped.

## 2 Discrete model

We use a conceptually straightforward exclusion process on a hexagonal lattice to simulate cell migration and cell proliferation (Jin et al., 2017). We apply this model to simulate *in vitro* cell proliferation assays, such as the experimental image in Figure 1. For this kind of proliferation assay, a standard way to estimate the proliferation rate is to count individual cells, and to use this information to construct time evolution of the average density profile (Jin et al., 2016). With this averaged density profile, we could calibrate the solution of the classical logistic equation to provide an estimate of the proliferation rate (Tremel et al. 2009). However, this standard approach neglects any variation in the proliferation rate. Therefore, here we use our model and take a different approach.

In our model, agents represent individual cells, and these agents are placed uniformly, at a specified initial density, on a hexagonal lattice. We use a lattice of size *I* × *J* lattice sites, with lattice spacing 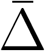. Here, 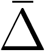 can be thought of as a typical cell diameter, such as 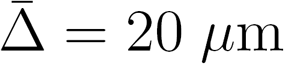. To simulate crowding effects, each lattice site can be occupied by, at most, one agent. Each lattice site is indexed, s = (*i,j*), where *i*, *j* ∈ ℤ^+^, and each site is associated with a unique Cartesian coordinate. The total population of agents is composed of *N* ≥ 1, potentially distinct, subpopulations. Agents in each subpopulation are characterised by a potentially distinct migration probability per time step, 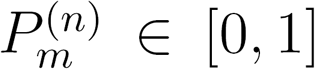 for *n* = 1, 2,…,*N*. Furthermore, agents in each subpopulation are characterised by a potentially distinct proliferation probability per time step, 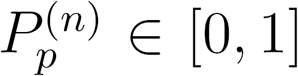 for *n* = 1, 2,…,*N*. The total number of agents at time *t* is 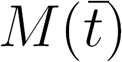.

Cell migration and proliferation are modelled using a random sequential random update method (Chowdhury et al., 2005). To advance the discrete model from time 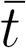 to time 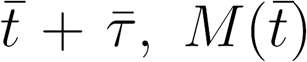 agents are selected at random, one at a time, with replacement. The selected agent attempts to move to one of the six nearest neighbour sites with probability 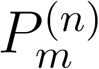. The attempted migration event will be successful if the randomly chosen nearest neighbour target site is vacant. After 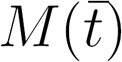 potential migration events have been attempted, another 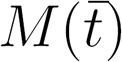 agents are selected at random, one at a time, with replacement. The selected agent will attempt to place a daughter agent on one of the six nearest neighbour sites with probability 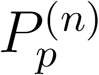. The attempted proliferation will only be successful if the randomly chosen nearest neighbour site is vacant. In the event that the potential proliferation event is successful, we make the simplest assumption that the daughter agent belongs to the same subpopulation as the mother agent. Once these potential motility and proliferation events have been attempted, we update 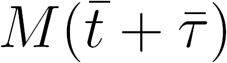.

Using a random sequential update algorithm means that during each time step some agents may attempt to migrate or proliferate multiple times, whereas other agents may not attempt to migrate or proliferate at all. However, when we simulate over a large number of time steps, on average each agent in the population will attempt to migrate and proliferate once per time step. The random sequential update algorithm is conceptually straightforward, easy to implement, and is known to provide a good approximation to the dynamics of these kinds of simulations where we consider populations of motile and proliferative agents (Treloar et al., 2014).

## 3 Continuum limit description

The continuum limit description of the discrete model can be derived using standard averaging arguments and a mean field approximation. These kinds of arguments and approximations are widely invoked throughout the mathematical biology literature where continuum limit descriptions are derived from underlying stochastic models (e.g. Callaghan et al., 2006; Deroulers et al. 2009; Dyson and Baker, 2015; Plank and Simpson, 2012). The mean field assumption involves treating the occupancy status of lattice sites as being independent. While this assumption is questionable for any particular single realisation of the stochastic model, it turns out to be remarkably accurate when we consider an ensemble of stochastic simulations, as we will show later. Furthermore, this approximation is known to be accurate for a range of problems involving interactions between multiple subpopulations of agents (Simpson et al, 2009), and for other problems involving single populations of motile and proliferative agents (Simpson et al. 2010).

We denote the probability of finding an agent from subpopulation n at site s = (*i,j*) as 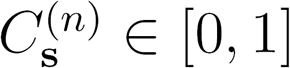, for *n* = 1, 2,…, *N*. This probability can be thought of as corresponding to averaging the occupancy of site s over many identically prepared realisations of the stochastic model. Therefore, the probability of site s being vacant is 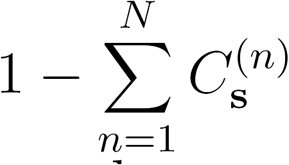. In the discrete model, migration events can act to either increase or decrease the occupancy of site s, whereas proliferation events can only act to increase the occupancy of site s. Accounting for these possibilities, the change in average occupancy at site s for agents from subpopulation *n*, from time 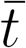 to time 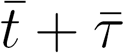 can be written as,

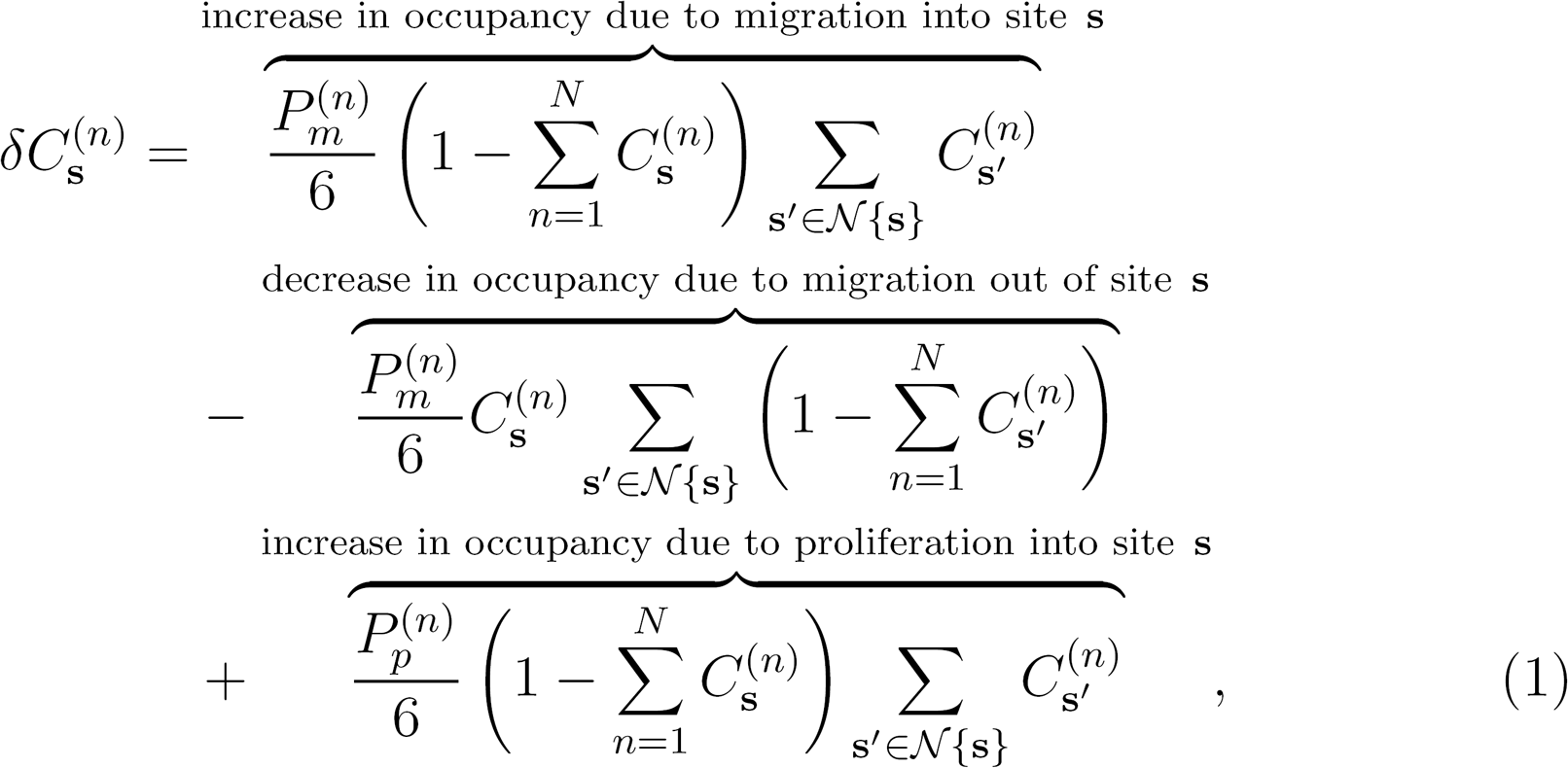

where 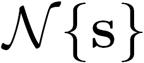 denotes the set of six nearest-neighbour sites around site **s**. In Equation (1) we implicitly make the standard assumption that the average occupancy of each lattice site is independent. This is the mean field assumption. We expand each term in Equation (1) about site s using Taylor series, and neglect terms of 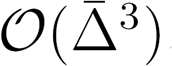. This process leads to the cancellation of many terms.Here, we omit showing these intermediate steps as they have been outlined for similar models in our previous work (Jin et al., 2016; Simpson et al., 2009; Simpson et al., 2010). Dividing both sides of the resulting expression by 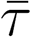, and taking the limit as 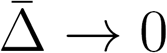 and 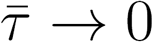 jointly, with 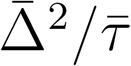 held constant we obtain,

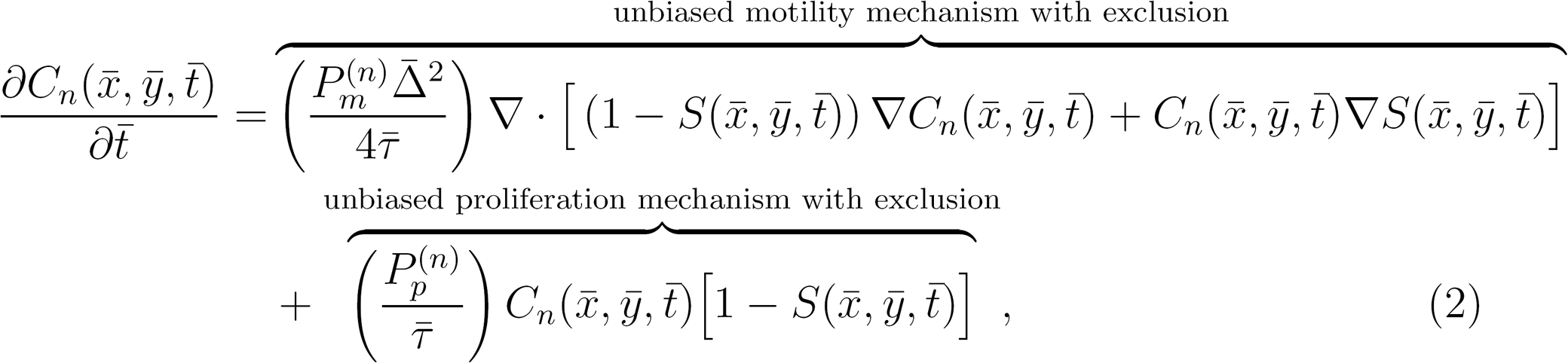

for *n* = 1, 2,…, *N*. 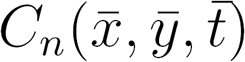 is the density of *n*^th^ subpopulation, and 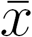 and 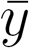 are the horizontal and vertical coordinates, respectively. The total population density is given by 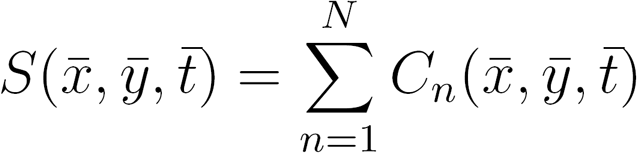.

In cell proliferation experiments, cells are placed uniformly on a two-dimensional substrate (Browning et al. 2018; Jin et al., 2017). Therefore, this kind of initialisation means that there are, on average, no spatial gradients in cell density provided we view the experiment at a sufficiently large spatial scale, such as in Figure 1. Under these conditions Equation (2) simplifies to

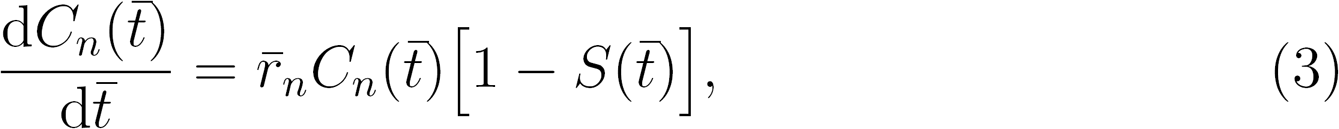

where 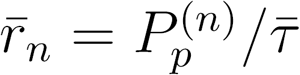, for *n* = 1, 2, 3,…, *N*. The total cell density 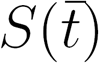 then can be obtained by summing over the governing equations of *N* subpopulations to give,

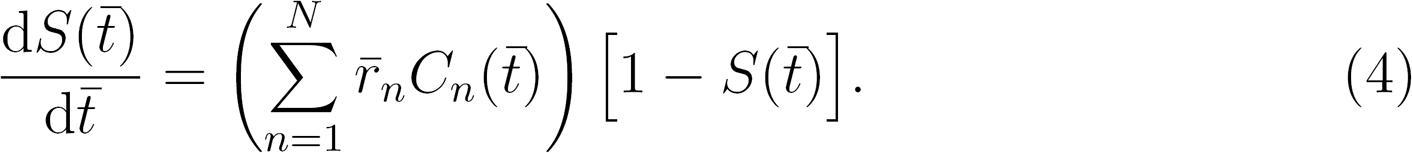

In this work we always deal with initial conditions without any spatial gradients, which corresponds to the experimental images shown in Figure 1. This implies that we are working with a system of ordinary differential equations (ODEs) instead of a system of partial differential equations. If we were to consider a different initial condition, such as a scratch assay or a barrier assay where there experiments are intentionally initialised with some spatial gradients present, then we would have to work with Equation (2) instead of Equation (3).

Without loss of generality, when we apply Equation (3) we adopt the convention that 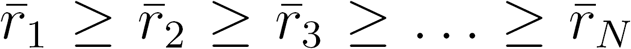, so that 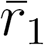 is the proliferation rate of the fastest-proliferating subpopulation, 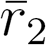 is the proliferation rate of the second fastest-proliferating subpopulation, and so on. We note that in the special case where we consider all the proliferation rates to be equal, 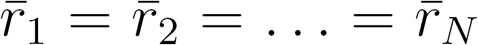, we are dealing with a homogeneous population with a constant proliferation rate, 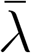. The continuum limit description simplifies to,

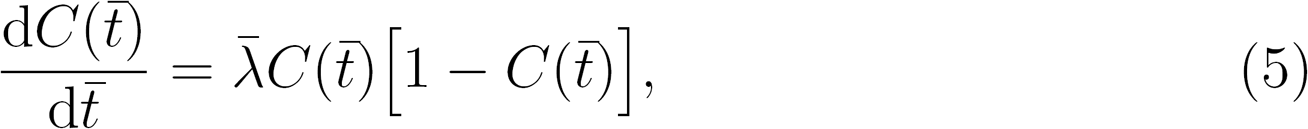

which is the classical logistic growth model (Murray, 2002), whose solution is given by,

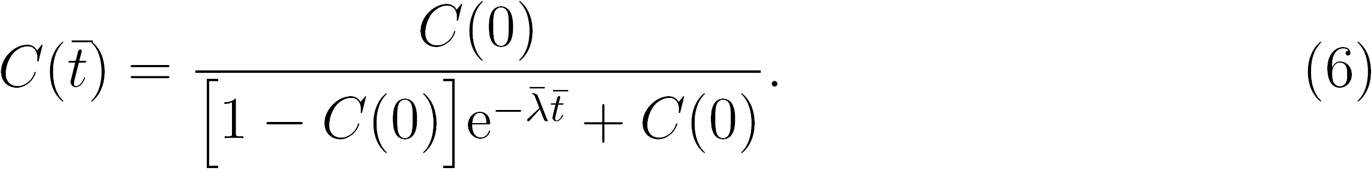

This exact solution is a sigmoid curve that monotonically increases from *C*(0), and approaches unity as 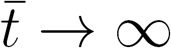, provided that *C*(0) < 1. Since our system of ODEs, given by Equation (3), simplifies to the classical logistic model when all the proliferation rates are identical, we refer to Equation (3) as the *extended logistic growth model.*

To simplify our work we nondimensionalise time using the fastest proliferation rate, 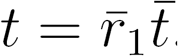. Therefore Equation (3) becomes

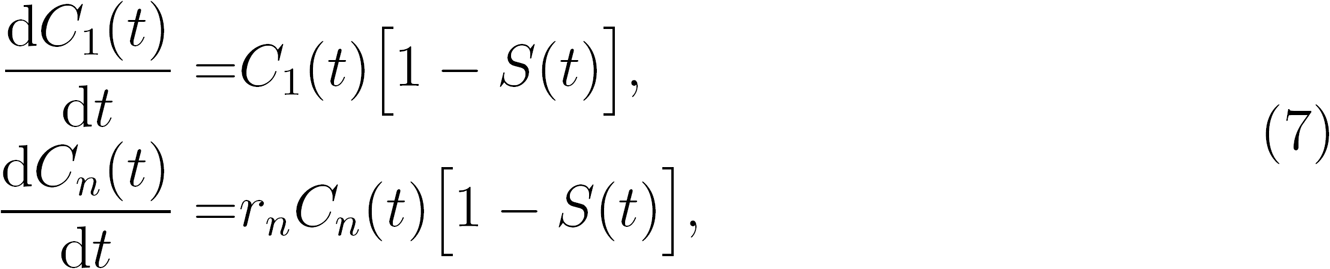

where 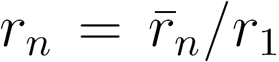, for *n* = 2, 3,…, *N*. Therefore, we now have a system of ODEs with non-dimensional proliferation rates of: unity, *r_2_,r_3_,…, *r*_N_*, with 1 ≥ *r_2_ ≥ r_3_ ≥…≥ *r*_N_*. This non-dimensionalisation allows us to compare the solutions of the model for different systems that are characterised by very different proliferation rates. In the non-dimensional format, Equation (4) be-comes

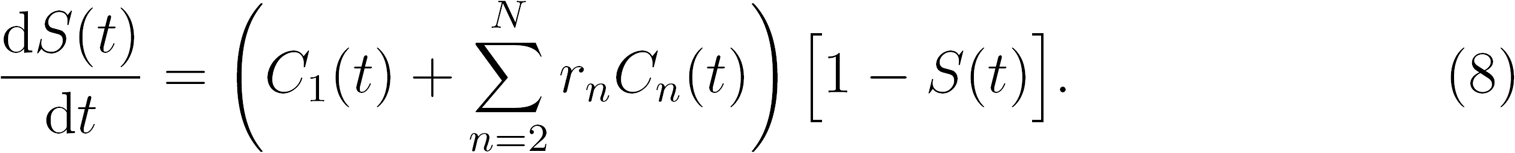

To be consistent, if we non-dimensionalise Equation (6) with 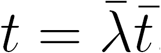 we obtain

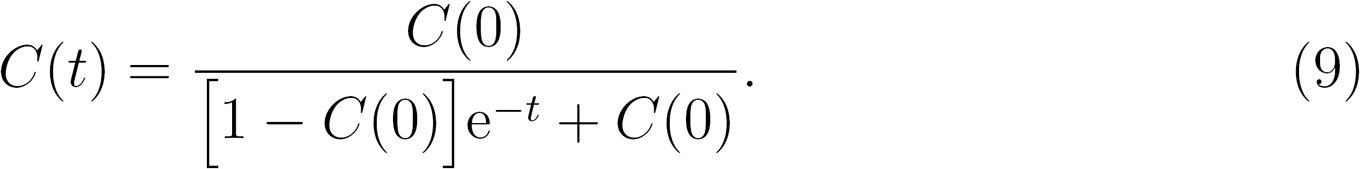

Unlike the classical logistic model, Equation (7) does not have an exact solution. Therefore, we present numerical solutions that are obtained using a backward Euler approximation. In all cases we use a constant time step of *δt* = 0.01, and Picard iteration with convergence tolerance ∈ =1 × 10^−5^. These choices of δt and ∈ are sufficient to produce grid-independent numerical solutions of the model.

## 4 Results

### 4.1 Continuum-discrete match

All discrete results are presented in a non-dimensional format, on a lattice with unit lattice spacing and with time steps of unit duration, Δ = τ = 1. Note that Δ and τ can be re-scaled to correspond to any particular choices of dimensional 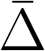 and 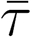. This means that we can re-scale any of these dimensionless simulations to correspond to a population of cells with arbitrary cell diameter, and arbitrary characteristic proliferation rate. Since we are focusing on the role of heterogeneity in cell proliferation, in all simulations we set 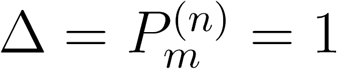, for *n* =1, 2,…, *N*, and 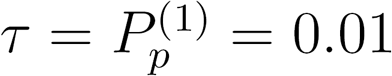 In addition, we choose *I* = 101 and *J* = 117, so that the size of the simulation domain is 100 × 100. Periodic boundary conditions are applied to all simulations.

To explore the role of heterogeneity in population growth, we first consider simulations involving up to three subpopulations: subpopulation 1 has the fastest proliferation rate; subpopulation 2 has an intermediate proliferation rate; and subpopulation 3 has the slowest proliferation rate. We first perform three different types of discrete simulations initialised with different combinations of these three subpopulations. Each simulation is initialised so that the total number of agents occupies just 10% of the total number of lattice sites. In the first simulation we consider a homogeneous population that is composed entirely of agents from subpopulation 1, *N* = 1. The second simulation involves a heterogeneous population that is composed of equal proportions of agents from subpopulations 1 and 2, *N* = 2. The third simulation involves a heterogeneous population that is composed of equal proportions of agents from subpopulations 1, 2 and 3, *N* = 3. Snapshots from the discrete simulations are shown in Figure 2. A qualitative comparison of these snapshots shows that the growth dynamics are very different in the homogeneous and heterogeneous populations. Both the dynamics of the overall total population, and the dynamics of the various subpopulations depends on the details of the heterogeneity present in the system.

**Fig.2.**
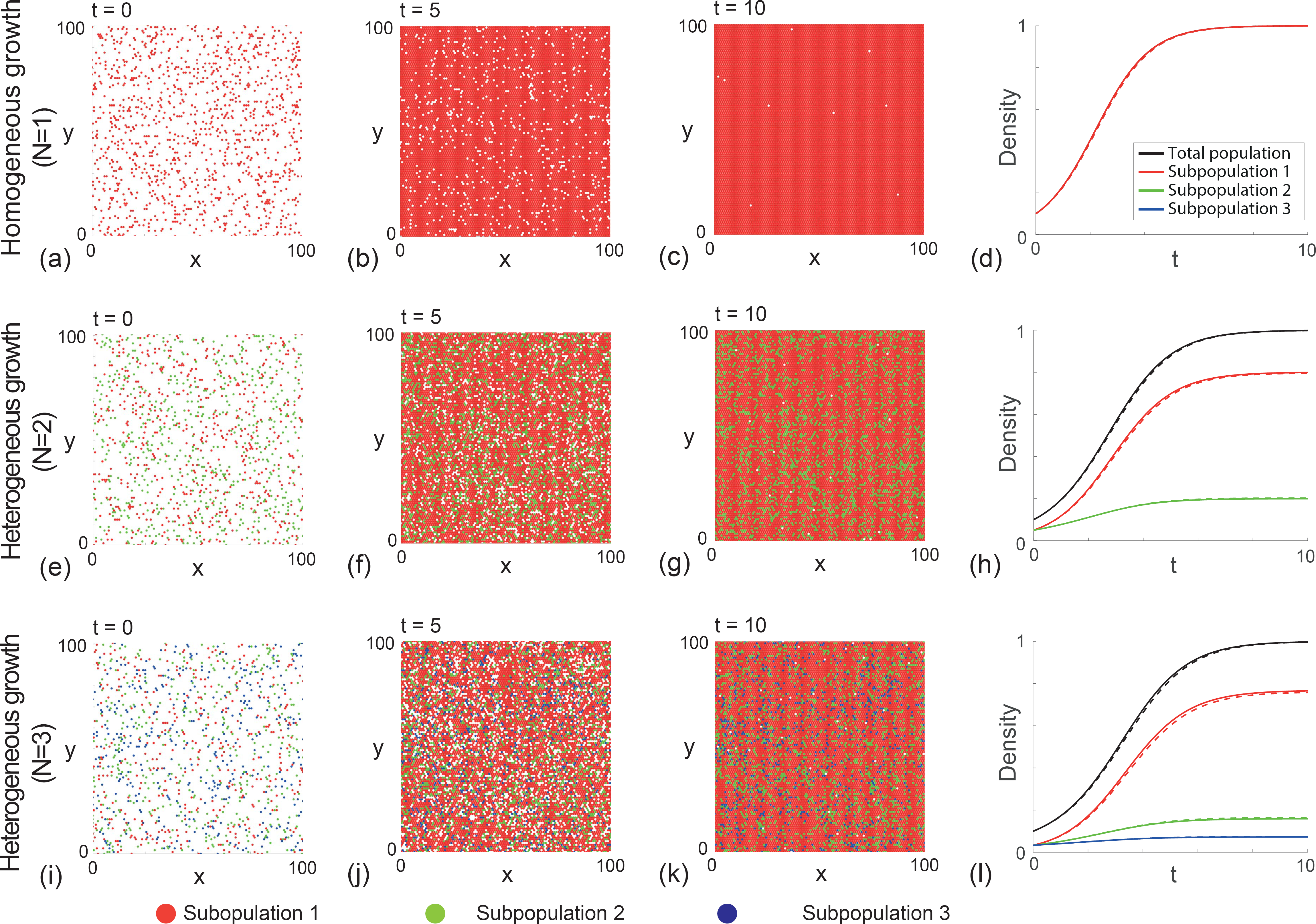
Discrete and continuum simulations of heterogeneous proliferation. Snapshots are shown at t = 0, 5, and 10 for: (*a*)-(*c*) a homogeneous population, *N* = 1; (*e*)-(*g*) a heterogeneous population, *N* = 2; and (*i*)-(*k*) a heterogeneous population, *N* = 3. In each case, 10% of lattice sites are initially randomly occupied, with equal proportions of the various subpopulations. (*d*), (*h*), (*l*) show the corresponding continuum-discrete matches. The solid lines are solutions of the extended logistic model, dashed lines are averaged discrete results, obtained by considering 50 identically prepared realisations. In all plots red represents the fastest-proliferating subpopulation (subpopulation 1); green represents the intermediate subpopulation (subpopulation 2); and blue represents the slowest-proliferating subpopulation (subpopulation 3). All simulations correspond to 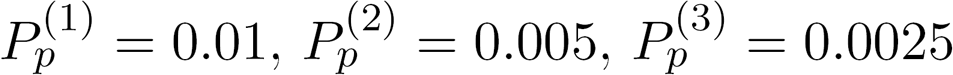, r_2_ = 0.5 and r_3_ = 0.25.

To quantify the population growth, we plot the time evolution of the total averaged agent density, and superimpose the corresponding solution of Equation (7). For the homogeneous population in Figure 2(d), the continuum limit simplifies to the classical logistic growth model, while for the heterogeneous populations in Figure 2(h) and (l), the extended logistic model applies. Overall we see that the quality of the match between the solution of the continuum model and averaged data from the discrete simulations is excellent. Therefore, this comparison indicates that the continuum limit description is a useful and accurate mathematical tool that can be used to study the proliferation of heterogeneous populations without relying on repeated stochastic simulations.

### 4.2 Comparison of the classical logistic growth model and the extended logistic model

As mentioned in the Introduction, although the classical logistic growth model is widely used when interpreting data from cell biology experiments (Cai et al., 2007; Jin et al., 2016; Maini et al., 2004; Sengers et al., 2007; Sheardown and Cheng, 1996; Sherratt and Murray 1990; Vo et al., 2015; Warne et al. 2017), this standard approach neglects any heterogeneity in cell proliferation rate. To provide insight into how well the classical logistic growth model is able to predict and describe the growth of heterogeneous populations, we now calibrate the solution of the classical logistic growth model in an attempt to match the solution of the extended logistic model which explicitly accounts for heterogeneous growth.

We consider two different initial conditions for a population that is composed of three different subpopulations, *N* = 3. Again, we refer to these subpopulations as subpopulations 1, 2, and 3. For both initial conditions we consider, we distribute the total population uniformly across the domain so that the initial total density is 10% of the carrying capacity density. In the first case we choose the initial condition so that the total population is initially composed of 75% of agents from subpopulation 1, 20% of agents from sub population 2, and 5% of agents from subpopulation 3, as shown in Figure 3(a). In the second case we choose the initial condition so that the total population is initially composed of 5% of agents from subpopulation 1, 20% of agents from subpopulation 2, and 75% of agents from subpopulation 3, as shown in Figure 3(b). These choices of initial condition mean that the first case is composed of a small proportion of relatively quiescent agents (di Fagagna et al., 2003), and the second case corresponds to a population that contains a small proportion of rapidly proliferating agents, such as is thought to be relevant to cancer progression (Davis et al., 2017). The solution of the extended logistic growth model for these two scenarios of heterogeneous growth are shown in Figure 3(b) and (d), respectively.

**Fig.3.**
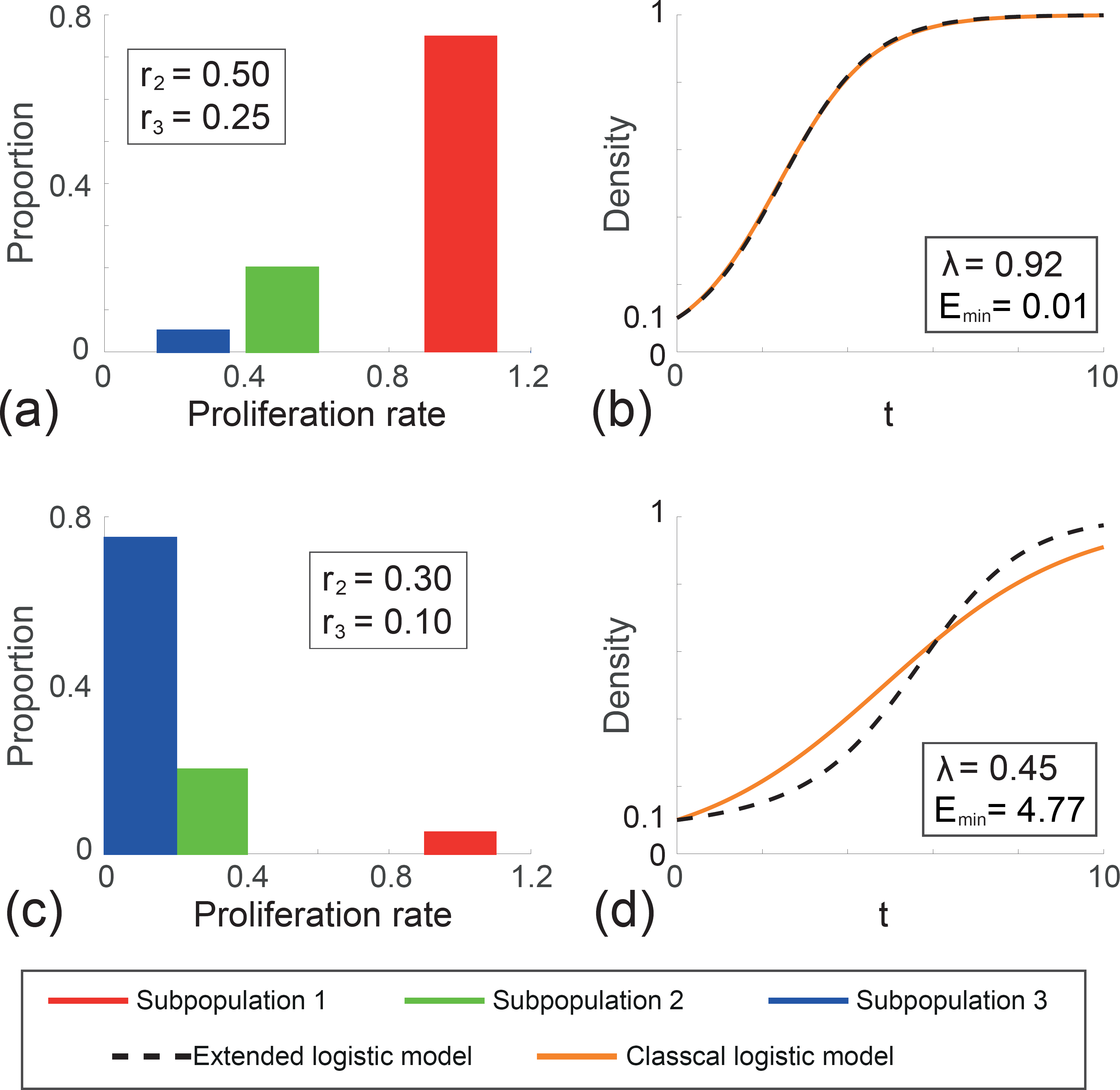
Comparison of the classical logistic growth model to the extended model for heterogenous growth. (*a*) and (*c*) Initial distribution of proliferation rate for the two heterogenous populations. (*a*) corresponds to a population composed of a small proportion of relatively quiescent cells, and (*c*) corresponds to a population containing a small proportion of rapidly proliferating cells. In all cases, red represents the fastest-proliferating subpopulation (subpopulation 1); green represents the intermediate subpopulation (subpopulation 2); and blue represents the slowest-proliferating subpopulation (subpopulation 3). In each case, the time evolution of the total density from the extended model, together with the best-fit classical logistic growth curve, is plotted in (*b*) and (*d*).

Again, we recall that standard approaches to interpreting cell proliferation assays is to calibrate the solution of the classical logistic growth model to match the experimental data, and to provide an estimate of the proliferation rate (Cai et al., 2007; Jin et al., 2016; Maini et al., 2004; Sengers et al., 2007; Sheardown and Cheng, 1996; Sherratt and Murray 1990; Vo et al., 2015; Warne et al. 2017). This standard approach implicitly neglects the role of heterogeneity, so it is of interest for us to take the population growth curves for these heterogeneous populations in Figure 3(b) and (d), and to calibrate the solution of the classical logistic growth model to match this data. We calibrate the solution of the classical logistic growth model to the density data in Figure 3(b) and (d) using MATLAB’s *lsqcurvefit* function, and show the best match in Figure 3(b) and (d). This calibration provides an estimate of A that is associated with the best fit of the standard model to the total density data. We are also interested in understanding the quality of match between the best fit solution of the classical logistic growth model and the heterogeneous density data. To quantify the quality of match we use a least-squares measure,

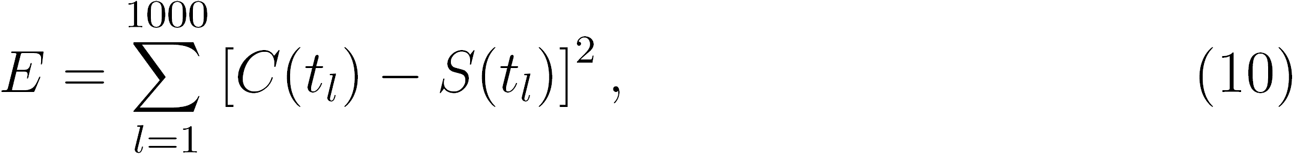

where *C*(*t*) is the best fit solution of the classical logistic growth model, and *S*(*t*) is the total density associated with the extended logistic growth model. Here we measure *E* in the interval 0 ≤ t ≤ 10, by evaluating both *C*(*t*) and *S*(*t*) at 1000 equally-spaced time points, *t_l_* for *t_l_* = 0, 0.01, 0.02,…, 10. We denote the minimum least-squares error as *E*_min_.

A simple visual comparison of the best fit classical logistic growth model and the solution of the extended model in Figure 3(*b*) shows that the standard approach of neglecting heterogeneity leads to an excellent match. However, results in Figure 3(d) indicate that the best fit classical logistic model matches the heterogeneous growth curve poorly. This qualitative assessment of the quality of match is confirmed quantitatively by our estimates of E, as reported in Figure 3. Overall, we see that the standard approach of neglecting heterogeneity can sometimes lead to reasonable outcomes, whereas in other cases the neglect of heterogeneity is unsatisfactory. We anticipate that the initial distribution of proliferation rates across the initial subpopulations plays an important role in determining the suitability of this standard approach. We now investigate this question further by applying our extended model to some data from the literature

## 5 Case study

We will now compare the performance of the classical logistic growth model and the extended logistic growth model by simulating proliferation assays in which the distributions of heterogeneous proliferation rates are taken from recent experimental measurements. In particular, we work with data from two human melanoma cell lines (Haass et al., 2014): (*i*) the 1205Lu cell line, which we refer to as cell line 1, and (ii) the WM983C cell line, which we refer to as cell line 2. The data we use to characterise the distribution of proliferation rates comes directly from Haass et al. (2014) where they use a specialised fluorescent technique to characterise the cell cycle of individual melanoma cells. Data from Haass et al. (2014) reports the duration of time spent in the S/G2/M phase of the cell cycle for groups of at least 20 individual cells from multiple melanoma cell lines. Since the S/G2/M cell cycle phase is closely related to the process of cell division, we treat the heterogeneity in these measurements as being representative of the heterogeneity present in the entire cell cycle.

Haass’ data reports the duration of time that at least 20 individual cells spend in the S/G2/M cell cycle phase (Haass et al., 2014). We group these individual measurements of duration into three subgroups. We choose the subgroups so that each column in the histogram of this data has approximately the same width, as shown in Figure 4(a)-(b). Since Haass’ data is reported in terms of a duration of time spent in the cell cycle, we convert these durations into rates by dividing log_e_(2) by the reported durations. Here, the log_e_(2) term comes from making the simple assumption that cells are growing exponentially. The details of the raw experimental data are given in the Supplementary Material document. Presenting the dimensional rates in Figure 4(c)-(d) indicates that the distribution of proliferation rate in cell line 1 is approximately symmetric, whereas the distribution of rates for cell line 2 is positively skewed.

**Fig.4.**
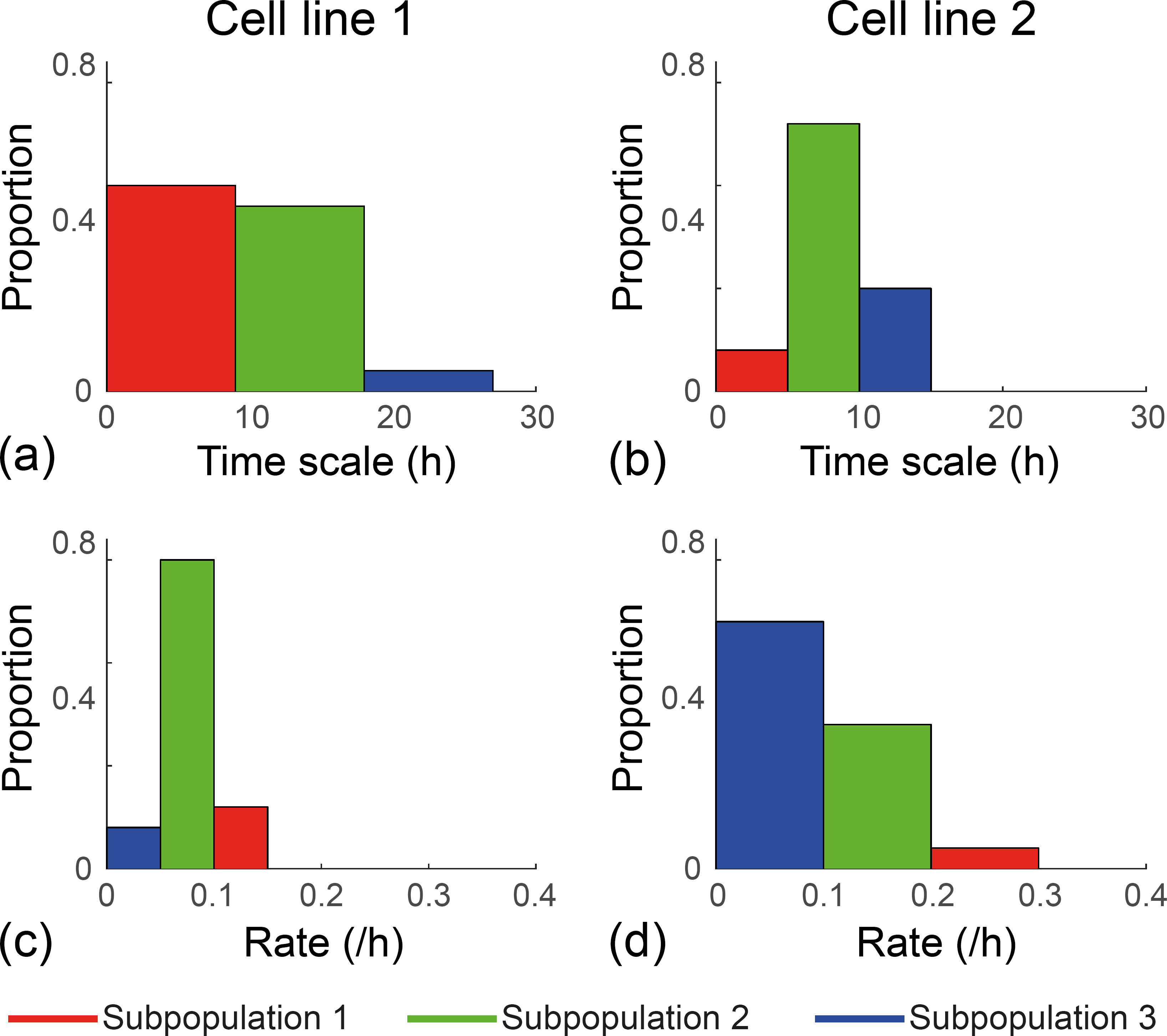
Experimental data from Haass et al. (2014). (*a*)-(*b*) Histograms showing the distribution of the time scale associated with the S/G2/M cell cycle for cell lines 1 and 2, respectively. (*c*)-(*d*) Histograms showing the distribution of rates associates with the S/G2/M cell cycle for cell lines 1 and 2, respectively. The time scales are converted to rates by dividing log_e_(2) by the time scale. For both cell lines: red indicates the fastest-proliferating subpopulation (subpopulation 1); green indicates the intermediate subpopulation (subpopulation 2); and blue indicates the slowest-proliferating subpopulation (subpopulation 3).

To apply our models using the heterogeneous proliferation rate data in Figure 4, we non-dimensionalise the proliferation rate data by dividing each rate by the fastest-proliferation rate for each cell line. This data, presented as histograms in Figure 5(a)-(e), shows the distribution of non-dimensional proliferation rates for both cell lines. We now use these histograms to specify both the initial proliferation rates, and the initial distribution of the three subpopulations in the discrete model and the corresponding extended logistic continuum model.

**Fig.5.**
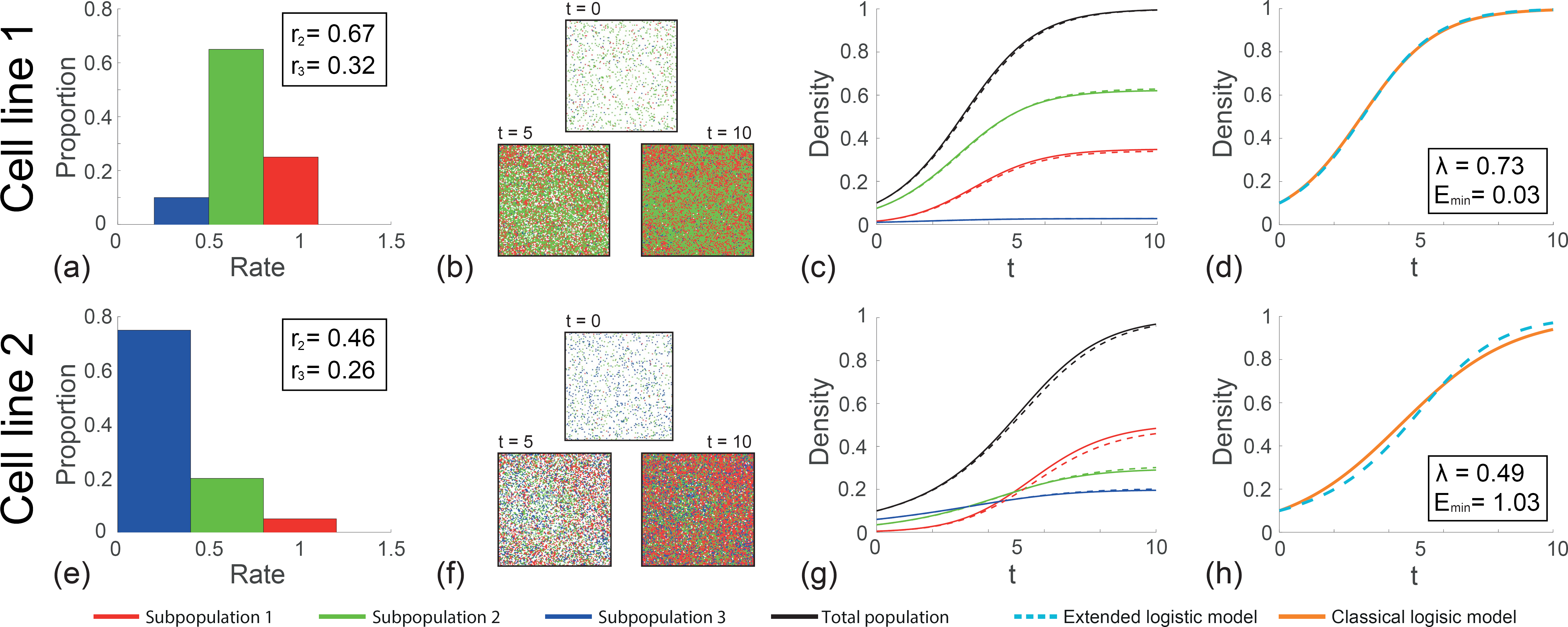
Comparison of the classical logistic growth model to the extended model for experimental cell lines 1 and 2. (a)and (e) Initial distribution of proliferation rate for cell lines 1 and 2, respectively. (b) and (f) Snapshots of simulations at t = 0, 5, and 10 for both experimental cell lines. (c) and (g) The continuum-discrete match of the time evolution of the densities for experimental cell lines 1 and 2, respectively. Solid lines represent the numerical solutions of the extended model, and the dashed lines represent the averaged simulation data over 50 identically prepared realisations. (d) and (h) Calibrated solution of the classical logistic growth model superimposed with the total density profile computed using the extended model. For both of experimental cell lines, red represents the fastest-proliferating subpopulation (subpopulation 1); green represents the intermediate subpopulation (subpopulation 2); and blue represents the slowest-proliferating subpopulation (subpopulation 3). For cell line 1, 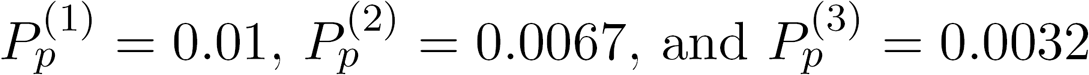. For cell line 2, 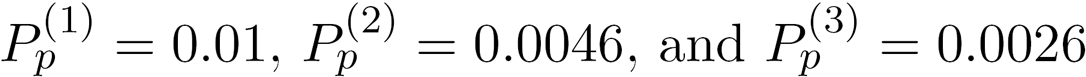.

We first model the heterogeneous population growth for cell line 1 and 2 using the discrete model. For each cell line, agents are initially distributed uniformly on the lattice so that 10% of lattice sites are occupied. We are also careful to ensure that the initial proportions of subpopulation 1, 2 and 3 correspond to the proportions of the three subpopulations in the histograms in Figure 5(a) and (e). Snapshots of the growing cell populations for both cell lines are given in Figure 5(b) and (f). These snapshots immediately reveal some interesting features. For cell line 1, the distribution of the three subpopulations remains similar over time, as there appears to be roughly equal proportions of red, green and blue agents at *t* =10 as there are initially, at *t* = 0. However, we observe very different behaviour for cell line 2, as the relative abundance of the three subpopulations changes dramatically over time. For example, at *t* = 0, we see that subpopulation 1 is the least abundant subpopulation. However, by the end of the growth process, at *t* = 10, subpopulation 1 is the most abundant subpopulation. These qualitative trends are also clear in Figure 5(c) and (g) where we compare averaged discrete data from repeated simulations of the stochastic model and the solution of the corresponding continuum model. In addition to quantifying the behaviour we see in the discrete snapshots, the solution of the continuum model in Figure 5(c) and (g) confirm that the continuum model is an accurate approximation of the discrete model.

To examine the implications of taking a standard approach and neglecting the role of heterogeneity, we also calibrate the solution of the classical logistic model to the total density data in Figure 5(c) and (g). Following the same approach described in Section 4.2, results in Figure 5(d) and (h) show the evolution of the total cell density profile superimposed with the best-fit classical logistic growth curves for cell line 1 and 2, respectively. Interestingly, the quality of match between the classical logistic growth model and the heterogenous population growth curve is relatively good for cell line 1, whereas the quality of match for cell line 2 is poor.

These results show that the consequences of neglecting the role of heterogeneity is subtle. In particular, under some circumstances it is possible to accurately predict the growth of a heterogeneous cell population using the classical logistic growth model, whereas in other circumstances the classical logistic growth model provides a poor match.

## 6 Analytical insight for two subpopulations, *N* = 2

To support our numerical solutions of the continuum model developed in Section 3, we now provide some simple analysis. This analysis provides both math-ematical insight into the extended logistic growth model, as well as providing biological insight into the effects of heterogeneity. For brevity we concentrate on the case in which there are two subpopulations present, with densities *C*_1_(*t*) and *C*_2_(*t*). In this case the extended model, Equation (7), simplifies to

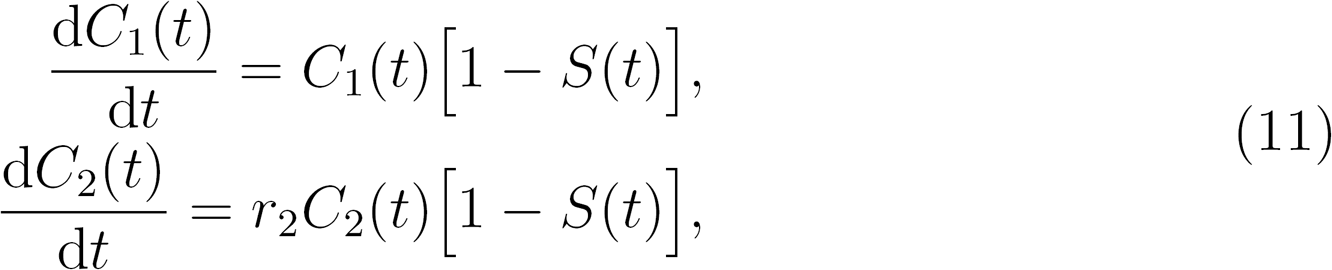

where *r*_2_ ≤ 1. The governing equation for the evolution of the total density is

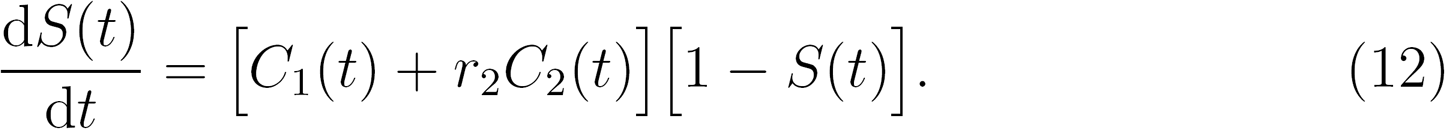

The solutions for both *C*_1_(*t*) and *C*_2_(*t*) are sigmoid curves that monotonically increase from the initial densities, *C*_1_(0) and *C*_2_(0), provided that *C*_1_(0) + *C*_2_(0) < 1. In the long time limit the solution of Equation (11) reaches a steady state solution, where *S*(*t*) → 1 as *t* → ∞. To analyse this long time behaviour we denote the steady state densities as,

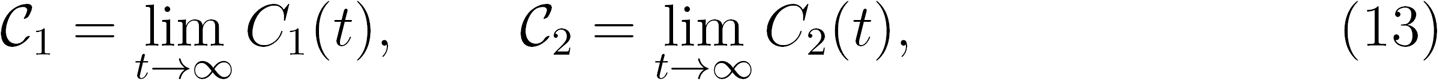

so that we have *C_1_* + *C_2_* = 1.

### 6.1 Exact steady state concentrations

It is not immediately clear what the steady state values 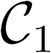 and 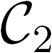 are from Equation (11) without first solving the transient model for the long time behaviour (Simpson et al., 2007). Furthermore, it is unclear how these steady state densities depend on the initial condition, *C*_1_(0) and *C*_2_(0), or on the proliferation rate *r*_2_. To provide insight into this question, we can solve for 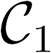 and 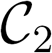 directly by first dividing one of the equations in (11) by the other, separating variables, and integrating to give

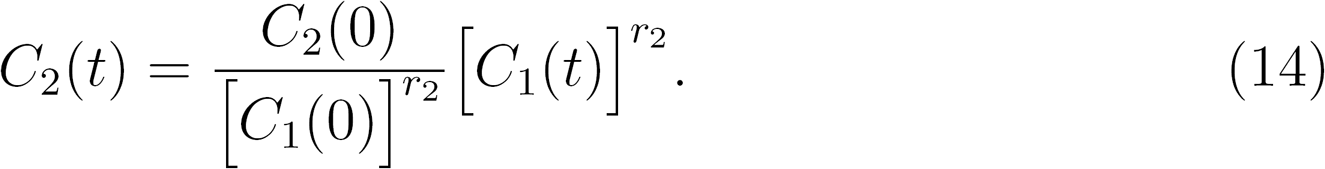

This relationship holds for all *t*. By substituting Equation (14) into Equation (11), we eliminate *C*_2_(*t*) to give

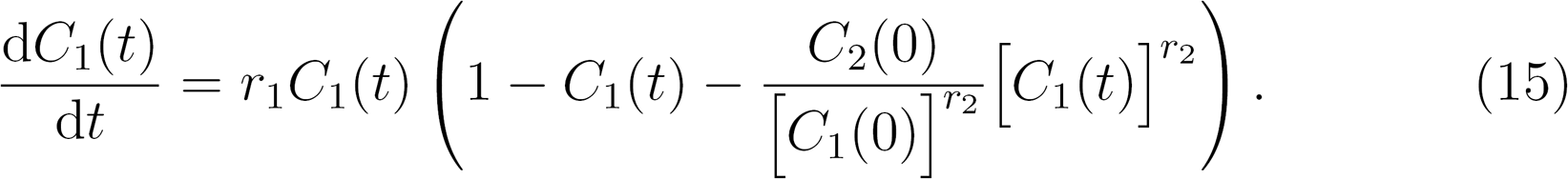

This equation is a now a direct analogue of the classical logistic growth model. For general values of *r*_2_ < 1, Equation (15) has no exact solution. However, in this form it is easy to read off the steady-state value by setting the time derivative to zero, resulting in

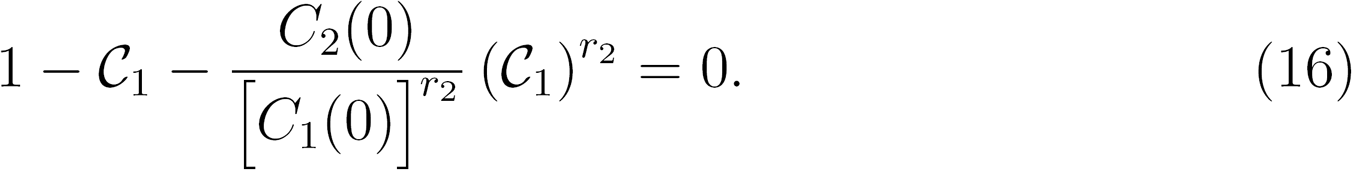

Equation (16) is a simple algebraic equation which can be solved using any iterative numerical method, such as MATLAB’s *fsolve* function. The value of 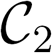 is then given by 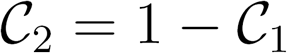.

Results in Figure 6(a)-(b) show two examples where we have used this approach to directly calculate 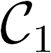 and 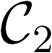. These predictions are superimposed on the associated transient solutions of Equation (11), showing that the direct method provides a simple and accurate way to calculate the long-time steady solution, without needing to use numerical integration to evaluate the long time limit of the transient solution.

**Fig.6.**
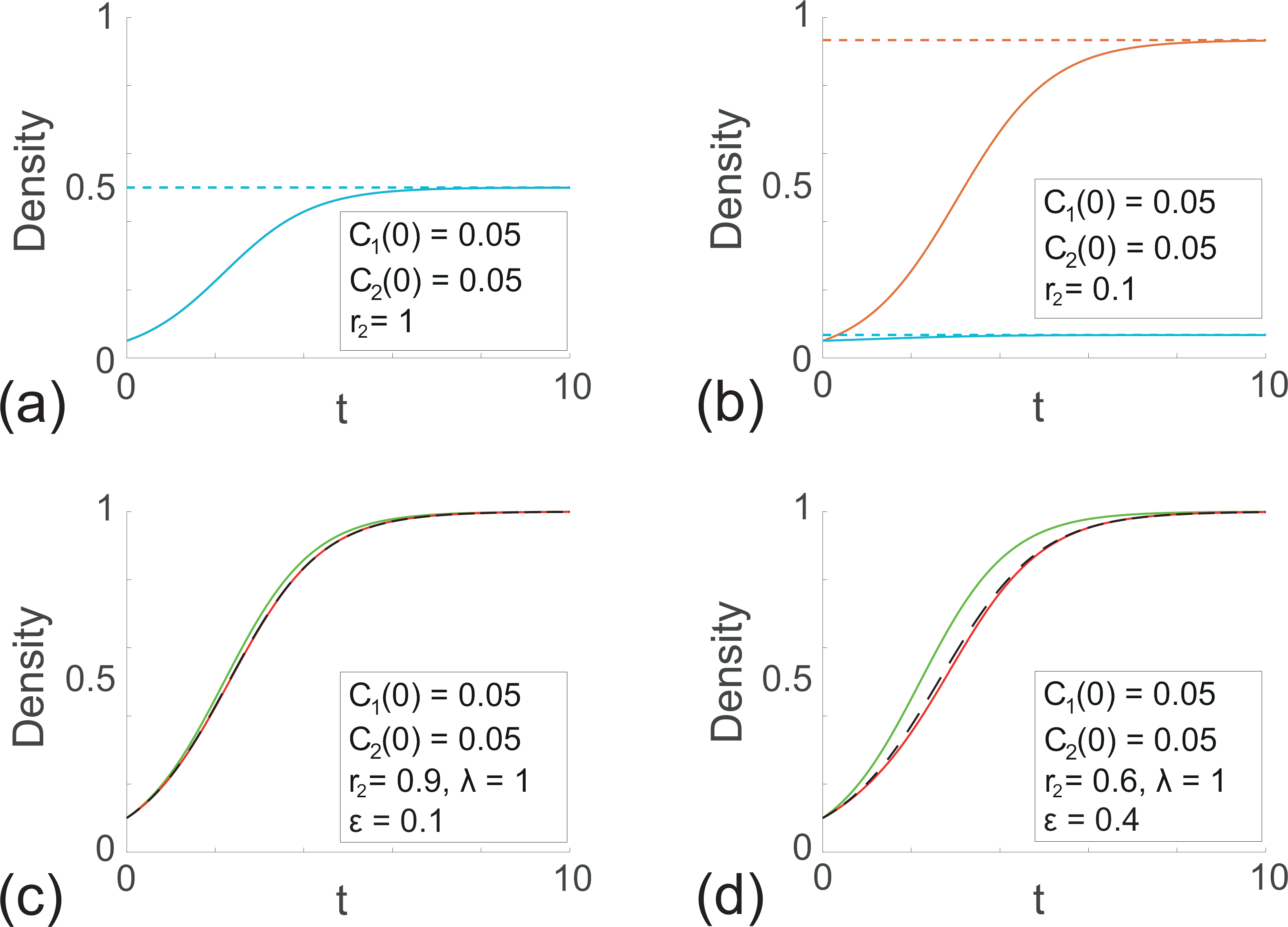
Analytical insight into the extended logistic model with two subpopulations, *N* = 2. (a)-(b) Steady state results (dashed) compared to the full transient numerical solutions (solid). Here, the first subpopulation is shown in orange, and the second subpopulation is shown in blue. (c)-(d) Comparison of the transient numerical solutions of total density (black) with the two-term perturbation solutions in the limit of small heterogeneity. The 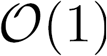 perturbation solution is plotted in green and the two-term 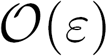 perturbation solution is plotted in red.

### 6.2 Approximate results for small heterogeneity

Although the extended logistic growth model given by Equation (11) does not have an exact solution, we can obtain approximate results in the limit of small heterogeneity. To explore this we consider *r*_2_ = 1 — ε, where ε ≪ 1, and propose the perturbation solution (Murray, 2012)

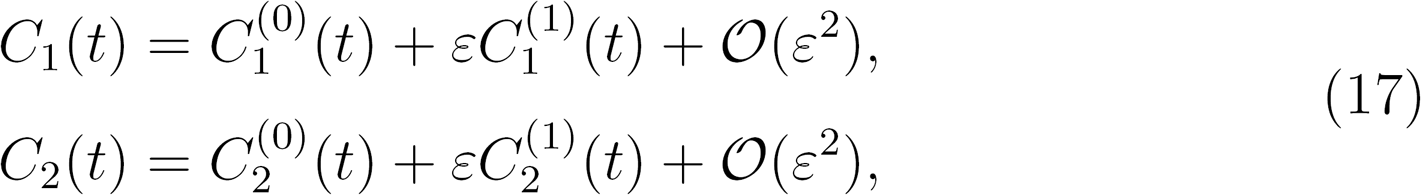

where the superscripts (0) and (1) represent the leading order and first correction terms, respectively. The asymptotic solution for the total population is obtained by summing over the solutions for the two subpopulations:

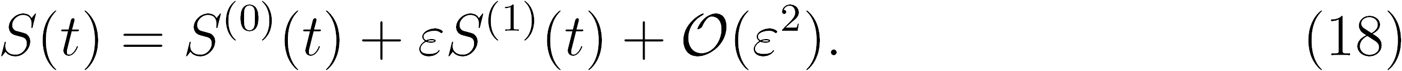

Substituting Equation (17) into the extended logistic model, given by Equation (11), gives the system

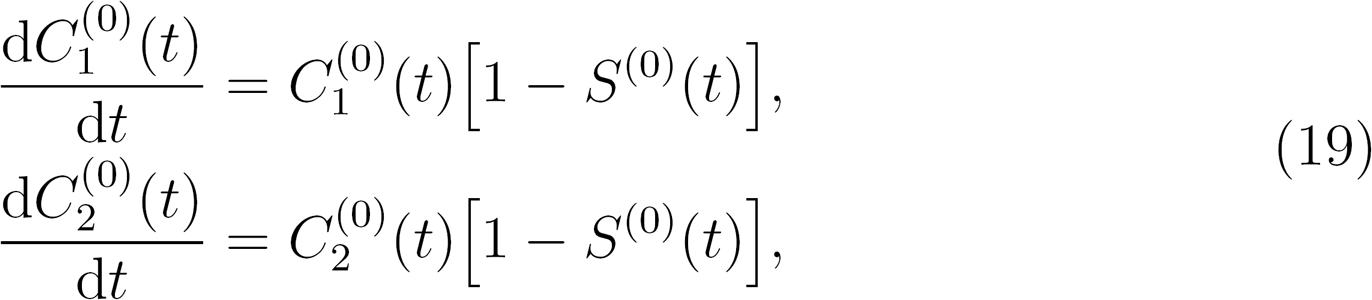

with 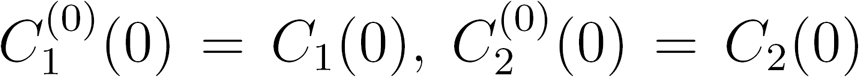. Correspondingly, the 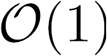 equation for the total population is

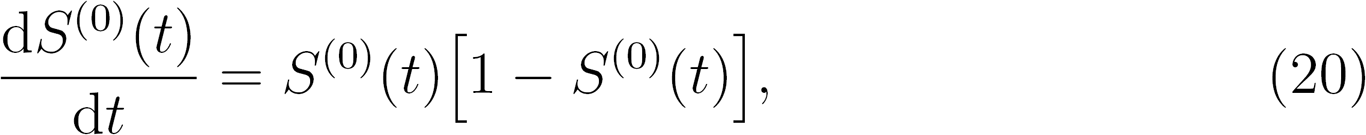

with S^(0)^(0) = S(0). Equation (20) is the standard logistic growth model with the explicit solution (Murray, 2002)

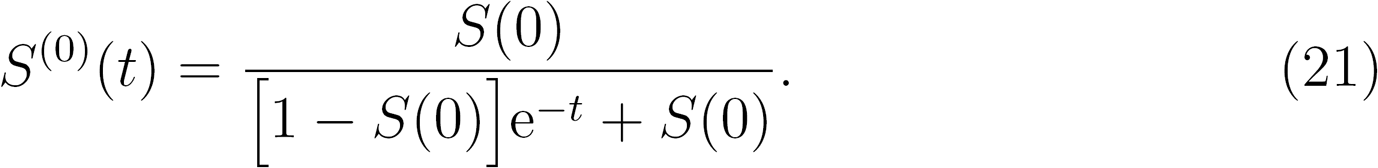

This is to be expected since our leading order problem holds for ∈ = 0, in which case both populations have the same proliferation rate, so effectively there is one population. Solving for the individual subpopulations gives the leading order solutions

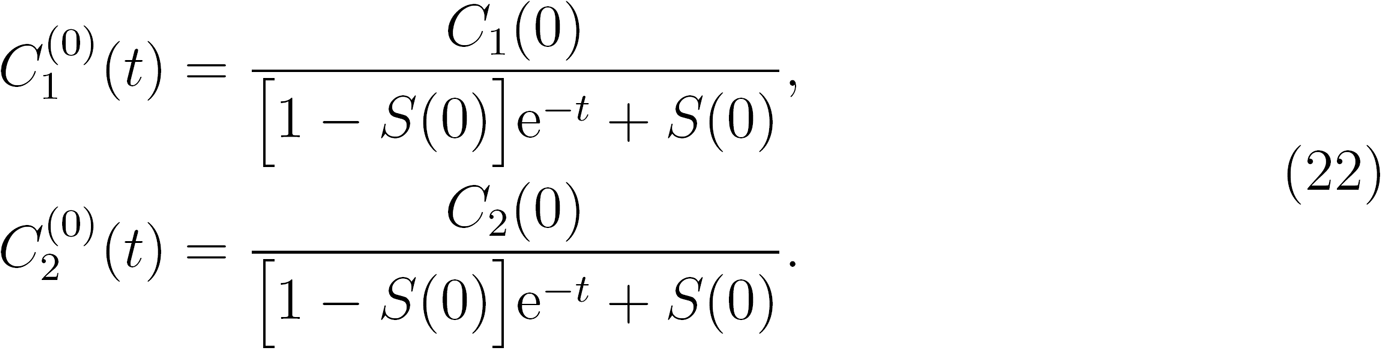

For ε ≪ 1, we proceed to solve for the correction terms. The governing equations for the individual populations are

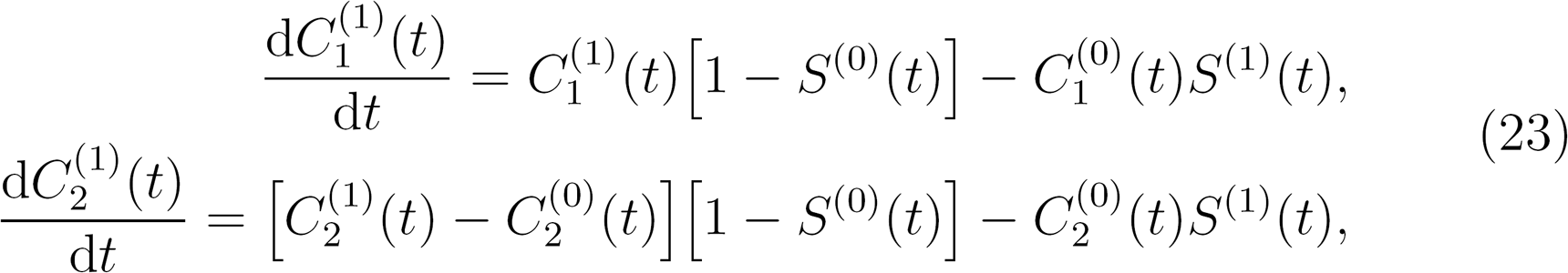

while the corresponding equation for the total density is

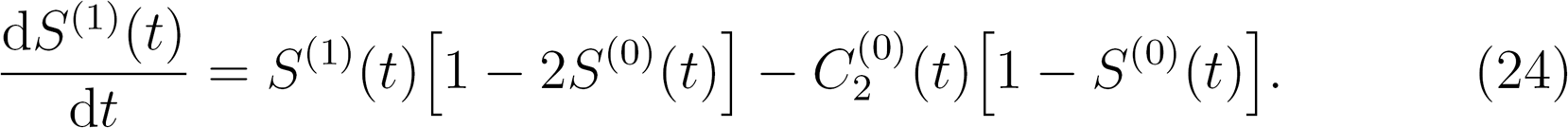

Equation (24) has an explicit solution

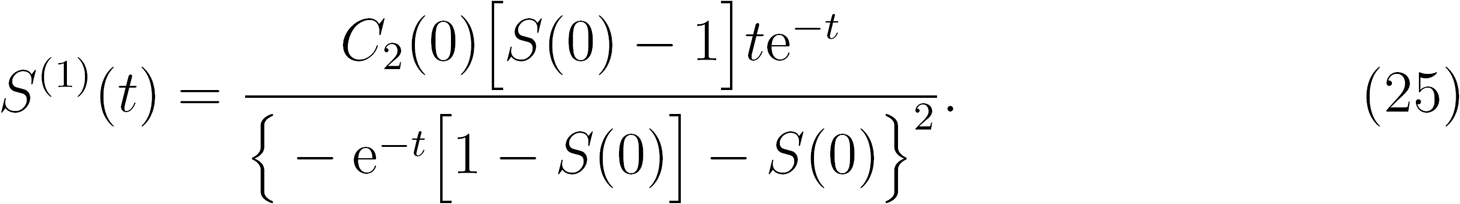

Neglecting higher order terms we obtain the two-term perturbation solution

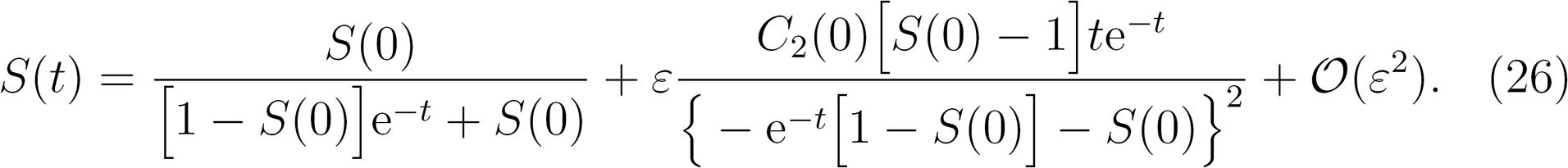

To demonstrate the effectiveness of this approximation, the 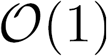 and 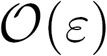 perturbation solutions for the total density are plotted in Figure 6(c)-(d). The corresponding full numerical solution is also presented. For a small amount of heterogeneity in the population (*ε* = 0.1, Figure 6(c)), the leading order term (solid green) provides a reasonably good approximation of the numerical solution (dashed black). However, for larger heterogeneity (*ε* = 0.4, Figure 6(d)), we see that the leading order term is no longer close to the numerical solution, and instead the full two-term perturbation solution Equation (26) (solid red) is required to provide a good approximation. These plots provide further evidence that when the population is almost homogenous, then the classical logistic model provides a good approximation. However, as heterogeneity becomes more pronounced, then our extended logistic growth model does a much better job at describing the dynamics. Furthermore, provided the heterogeneity is not too great, our two-term perturbation solution acts as a very good analytical approximation.

Results in Figure 6(c)-(d) correspond to one particular choice of initial condition, C_1_(0) and C_2_(0). Additional results (Supplementary Material, Figure S3) show that Equation (26) also provides a good approximation when we vary the initial condition, provided that ε is sufficiently small.

## 7 Conclusions

In this study we develop discrete and continuum models of cell migration and cell proliferation that allow us to explicitly investigate the role of heterogeneity, with a particular emphasis on the role of heterogeneity in the proliferation rate. Despite the fact that heterogeneity is commonly observed in cell populations, and is thought to play an important role in disease progression and tissue repair (Evan and Vousden, 2001; Haridas et al., 2017; Pavlath et al., 1998), standard mathematical models of cell migration and cell proliferation neglect to account for heterogeneity. Indeed, most standard mathematical models of cell proliferation simply treat the proliferation rate as a constant.

To explore the role of heterogeneity, we start by developing a discrete modelling framework to simulate cell migration and cell proliferation, modulated by crowding effects. The key point of the model is to deliberately introduce heterogeneity in the individuals within the population. The continuum limit description of the discrete model leads to a system of coupled, nonlinear ODEs. It is of interest to note that in the simplest case where the proliferation rates of each subpopulation are identical, the system of ODEs simplifies to the classical logistic growth model. Therefore, we call the new model the extended logistic growth model. Averaged data obtained from repeated simulations of the discrete model compare very well with the solution of the extended logistic growth model.

To explore the consequences of applying the logistic growth model and neglecting the role of heterogeneity, we perform a set of *in silico* experiments and generate density profiles describing the growth of a heterogeneous population of cells. We calibrate the solution of the classical logistic equation to match that data. Interestingly, while the classical logistic growth model can provide an accurate prediction of the growth of some kinds of heterogeneous populations, we also find that in some circumstances the classical approach can not make accurate predictions. We also generate *in silico* data by parameterising the extended logistic growth model with a set of heterogeneous proliferation rates from recent experimental measurements. Again, we find that the classical logistic model performs very well under some conditions, but it performs poorly for others. Overall, we find that when the heterogeneous population contains a small proportion of relatively fast proliferating cells, the classical logistic equation performs poorly. Therefore, we suggest that care ought to be exercised when modelling the growth of certain cell populations. For example, when modelling a population of cells that might involve mutations that act to increase the proliferation rate in a small subpopulation (e.g. Davis et al., 2017), the extended logistic growth model might be more accurate than the classical logistic equation.

Unlike the classical logistic equation, the extended logistic growth model does not have an exact solution. Therefore, in general, we have to rely on numerical solutions. However, we also show how to provide some further insight by obtaining analytical solutions in the case where there are just two subpopulations present, *N* = 2. We obtain exact expressions for the long-time steady state solution, and show that these exact expressions can be solved numerically to predict the steady state solution without using numerical integration to solve the full transient model. Furthermore, we also obtain approximate insight by constructing perturbation solutions in the limit that the degree of heterogeneity is small. The perturbation solutions are insightful since the 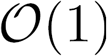 perturbation solution for the total cell density is the classical logistic equation. The 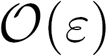 perturbation solution provides a correction term that is accurate even when we consider a relatively large degree of heterogeneity in the system.

Although we have focused here on the question of developing mathematical tools and mathematical insight into the role of heterogeneity in population dynamics associated with populations of cells, it seems likely that the ideas explored here will have consequences beyond the cell biology literature. For example, classical logistic models, with constant growth rates, are also commonly used in mathematical ecology (e.g. Chan and Kim, 2013), and there is also an awareness in the ecology literature that ecological populations do not always grow logistically (e.g. Taylor and Hastings, 2005). Therefore, perhaps some of the ideas developed here might also play a role in our understanding of ecological population dynamics. Another feature of our work is that we have focused exclusively on heterogeneity in cell proliferation rates. However, we note that there is also considerable interest in developing quantitative, predictive mathematical models which incorporate heterogeneity in cell migration (e.g. Read et al., 2016). Again, it seems likely that the kind of approach taken here would also be of interest in the context of exploring heterogeneity in cell migration. These open questions could be considered in future studies.

## Acknowledgments

This work is supported by the Australian Research Council (DP140100249, DP170100474). Wang Jin is supported by a QUT Vice Chancellor’s Research Fellowship. We thank two anonymous reviewers for their helpful comments.

